# Cortical folding patterns are encoded in the geometry of the unfolded neocortex

**DOI:** 10.64898/2026.06.05.730475

**Authors:** Roberto Toro, Katja Heuer, Nisrine Aflak, Victor Borrell, Clément Caporal, Camino de Juan Romero, Benoît Larrat, Antoine Legouhy, Erik Maikranz, Jean-François Mangin, Kevin Martinez-Anhom, Sébastien Meriaux, Isabel Reillo, Nicolas Traut

**Author notes:** Corresponding author: Roberto Toro.

## Abstract

Cortical folding patterns are conserved across individuals of gyrencephalic species and are closely related to cytoarchitectural organisation, connectivity, and function. Early morphogen gradients have been proposed as the molecular source of positional information encoding these patterns – a gyral molecular protomap – but the contribution of neocortical geometry to this encoding has not been examined. Here we show that the geometry of the unfolded ferret brain guides the adult folding pattern before any folding has occurred. Using high-resolution MRI surface reconstructions from postnatal day (P) 0 to adults, we demonstrate that newborn neocortical curvature predicts mature curvature maps, sulcal-gyral fate, and fold orientation. Pre-folding curvature correlates with expression of key patterning genes, and a mediation analysis indicates that geometry at P0 is the principal predictor of sulcal-gyral fate at P6. Mechanical simulations show that regions of high curvature act as autonomous anchors that organise the folding pattern. These results suggest that neocortical geometry constitutes a form of positional information that complements molecular patterns in regulating cortical organisation, and that a complete account of cortical development requires the integration of geometric and mechanical processes alongside molecular signalling.

## Introduction

The neocortices of gyrencephalic mammalian species fold in characteristic patterns. These patterns are stable across individuals, and present a remarkable relationship with neocortical cytoarchitecture, connectivity and functional organisation (Welker 1990, Heuer et al. 2025, Toro and Heuer 2025). These observations have led to the hypothesis that cortical patterns are genetically encoded (Akula et al. 2023, Singh et al. 2024, Moffat and Schuurmans 2024). One proposed mechanism is that early morphogen gradients convey positional information (TTkačik and Gregor 2021), as in the classic French Flag problem of Wolpert (Wolpert 1969, Sharpe 2019, Fig. 1a). In the specific case of the neocortex, a small number of morphogen gradients established early in development could provide each progenitor with a unique combination of concentrations, committing it to a region-specific fate, driving arealisation and guiding connectivity (Rakic 1988, Welker 1990, Van Essen 1997, O’Leary and Nakagawa 2002, Caronia-Brown et al. 2014). Mallamaci et al. (2000) showed that knockout of the homeogene *Emx2* in mice produced an anterior shift of cortical areas, and Fukuchi-Shimogori and Grove (2001) demonstrated that ectopic FGF8 in mouse embryos shifted area boundaries leading even to the partial duplication of the somatosensory barrel field. Such positional information signals could also regulate localised differential expansion, producing gyri and sulci (Ronan and Fletcher 2014, Lefèvre and Mangin 2010, Singh et al. 2024).

**Figure 1.**
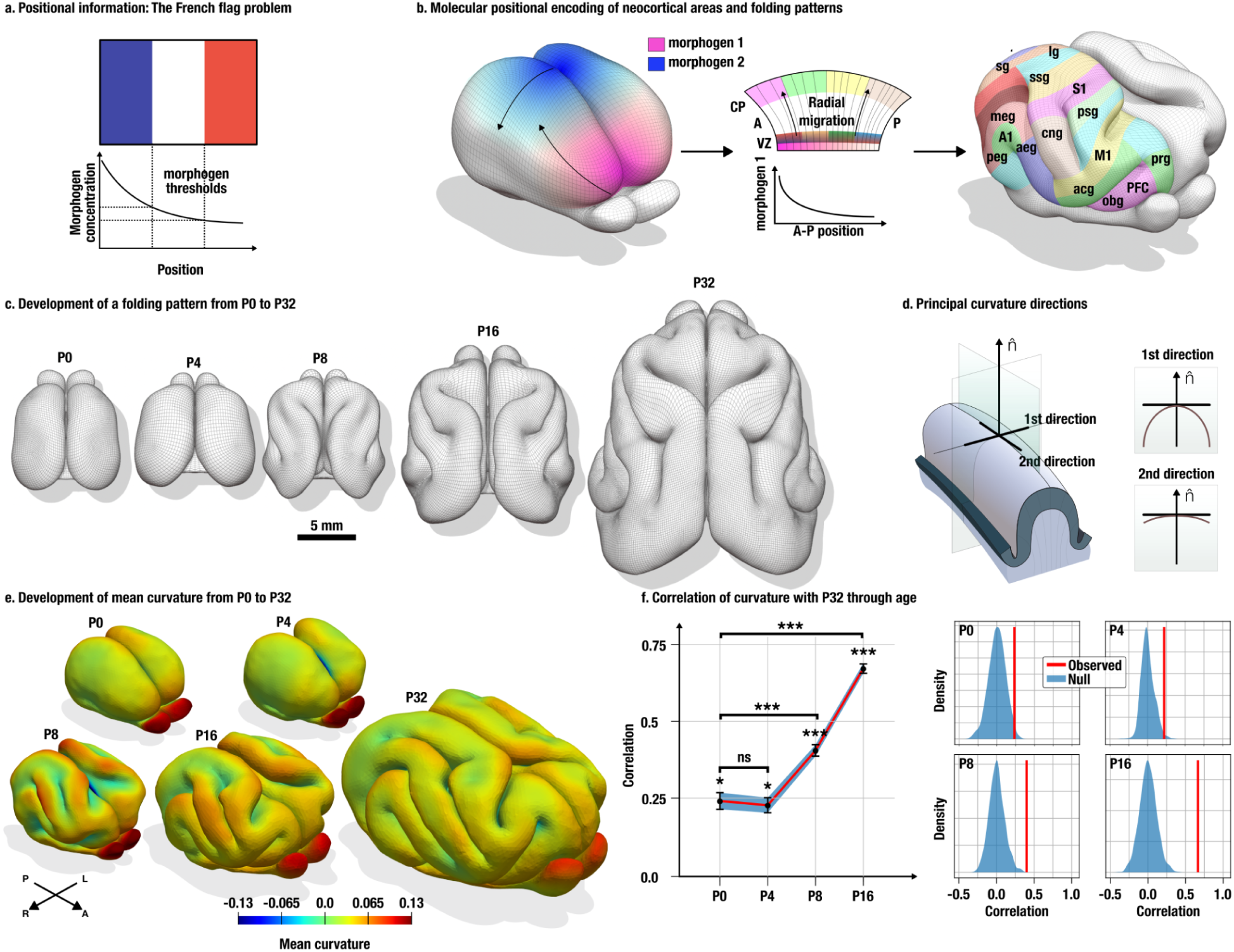
Correlation of the curvature map of the unfolded newborn ferret neocortex and after mature folding has been achieved. **a.** Positional information is an influential concept in developmental biology: a combination of morphogen gradients encodes position, which is subsequently interpreted as region-specific specialisation. In the classic example, a single gradient encodes the 3 regions of the French Flag (Wolpert 1969, Sharpe 2019). **b.** In the neocortex, positional information could similarly be encoded by signalling molecules secreted from a small number of patterning centres. This information underlies a primordial map of neocortical regions – the protomap (Rakic 1988, O’Leary 1989). Through radial migration, this protomap would be propagated from progenitor zones to the cortical plate, giving rise to cytoarchitectonic areas, connectivity patterns, and the folding pattern (Welker 1990, Van Essen 1997, Van Essen 2020). **c.** We propose that the early geometry of the neocortex provides an alternative – or complementary – positional signal capable of guiding the adult folding pattern. We study ferret neocortical development from birth (P0) to P32. At birth, the ferret brain is smooth. By the end of the first postnatal week, a folding pattern begins to emerge concomitantly with the specification of neocortical areas and cortico-cortical connectivity. By P32, the ferret brain has reached adult volume and ferrets open their eyes for the first time. **d.** The principal directions of curvature characterise the local geometry of a surface. At each point, a 1st principal direction (maximum curvature) and a 2nd principal direction (minimum curvature) can be defined. **e.** Mean curvature in the ferret neocortex is globally positive during the first postnatal week, with higher values near the medial wall. When folding develops, mean curvature maps become complex, combining positive values (gyri) and negative values (sulci). **f.** The mean curvature of the unfolded newborn neocortex correlates significantly with that of the adult brain. By the end of the first week, when folds start to appear, the prediction strengthens considerably. * p<0.05, ** p<0.01, *** p<0.001.

It has become increasingly clear, however, that cells can sense and interpret mechanical and geometric cues with a precision comparable to that of molecular signals (Pillai and Franze 2024). Tissue geometry is not only a consequence of growth but also an active regulator of it (Nelson et al. 2005) in a dynamic interplay between mechanics and molecular signalling (Heisenberg and Bellaïche 2013, Tzika et al. 2023). We hypothesised that the geometry of the developing neocortex provides geometric cues that complement molecular positional information, helping to specify which regions become gyri or sulci and to constrain their orientation.

To test this hypothesis, we studied neocortical folding in ferrets, an important animal model for brain development (Shinmyo et al. 2022, Garcia et al. 2025). Unlike humans, ferrets are born lissencephalic and with a very immature neocortex (Smart and McSherry 1986a, Smart and McSherry 1986b, Barnette et al. 2009, Sawada and Watanabe 2012). During the first postnatal weeks, the neocortex expands and a rich, stable folding pattern develops (Fig. 1c). By the end of the first month, when the ferret brain reaches the volume of an adult ferret, they open their eyes for the first time. From this perspective, the first postnatal month in ferret brain development recapitulates the last trimester of human gestation (Gilardi and Kalebic 2021).

We show that the curvature of the newborn ferret neocortex – one week before the onset of folding – correlates significantly with the curvature of the adult brain and predicts which regions become sulci or gyri. The principal curvature directions of the unfolded brain strongly constrain the orientation of mature folds. Reanalysis of published gene expression data shows that pre-folding curvature correlates with regional gene expression, and a mediation analysis indicates that curvature at P0 is the principal predictor of sulcal-gyral fate at P6. Finally, mechanical simulations demonstrate that regions of high initial curvature act as autonomous anchors that canalise the emerging folding pattern.

## Results

### The curvature pattern of the unfolded neocortex correlates with that of the folded neocortex

The mean curvature of the unfolded ferret neocortex at postnatal day 0 (P0) correlated significantly with the mean curvature of the folded brain at P32. We analysed a series of high-quality surface reconstructions from high-resolution ex-vivo MRI at P0, P4, P8, P16 and P32. The newborn ferret brain (P0, P4), has a characteristic bean-like shape. At every point of the surface, two principal curvature directions can be defined (Fig. 1d). The mean curvature – the average of these two curvatures – is an extrinsic measure of local curvature (a folded paper has non-zero mean curvature). The Gaussian curvature – the product of the two principal curvatures – is an intrinsic measurement of curvature, the type of curvature that cannot be removed by flattening (zero for a flat sheet or a cylinder; non zero for a sphere). We computed mean curvature maps at each stage (Fig. 1e). Mean curvature at P0 and P4 correlated significantly with that at P32 (Fig. 1f, P0: r=0.235 ± 0.027, p=0.014; P4: r=0.221 ± 0.025, p=0.024; standard errors by block jackknife, p-values by spin tests, Alexander-Bloch et al. 2018, Markello et al. 2022). As expected, correlations strengthened after P8, once the folding pattern was already apparent (P8: r=0.399±0.019; P16: r=0.666±0.015; p<0.001 for both).

### The curvature of the unfolded neocortex predicts mature sulcal-gyral fate

The mean curvature at P0 or P4 predicted whether a neocortical region would become a sulcus or a gyrus by P32. We applied cross-validated logistic regression using mean and Gaussian curvatures as predictors of sulcal-gyral fate at P32 (Fig. 2a). Sulcal and gyral regions were determined by thresholding the mean curvature maps. Prediction accuracy was evaluated as the area under the receiver operating characteristic (ROC AUC) curve (AUC; 0.5=chance; 1.0=perfect), and significance was estimated using spin tests. Before any folding, prediction was moderate but statistically significant (Fig. 2d, P0: AUC=0.65, p=0.034; P4: AUC=0.67 p=0.018). There was no significant difference in accuracy between P0 or P4. As expected, accuracy increased substantially after the onset of folding (Fig. 2e, P8: AUC=0.86, p≪1; P16: AUC=0.95, p≪1).

**Figure 2.**
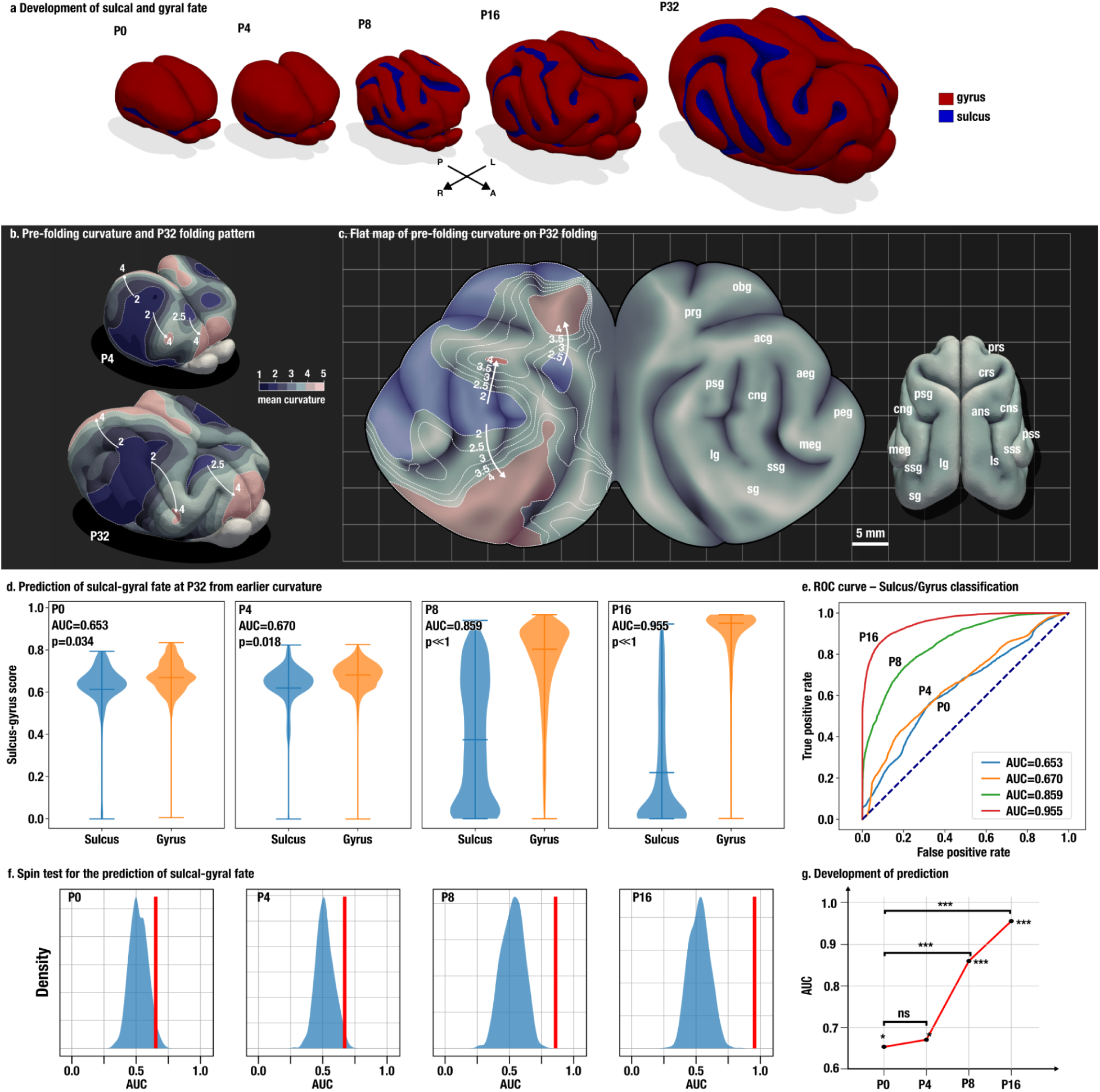
Prediction of sulcal-gyral fate. **a.** The ferret neocortex begins to fold by the end of the first postnatal week. Thresholding mean curvature maps provides a quantitative classification of sulcal (negative curvature, blue) and gyral (positive curvature, red) regions. **b.** Although mean curvature is positive everywhere in the newborn neocortex, it is not homogeneous: the hemispheric rim near the sagittal plane has notably higher curvature than the lateral surface. When this map is projected onto the folded P32 brain, high-curvature regions correspond predominantly to gyri and low-curvature regions to sulci. The direction of curvature gradients appears to anticipate the main axes of folding. **c.** Flat map of the P32 neocortex with pre-folding curvature projected to facilitate the inspection of this correspondence. **d.** Early curvature patterns predict sulcal-gyral fate with statistical significance before folding has begun; accuracy increases sharply at the onset of folding **e.** ROC curves for the prediction of sulcal-gyral fate. **f.** Significance estimated using spin tests. **g.** Development of prediction accuracy across postnatal stages.

An intuition for the mechanism driving the prediction can be obtained by inspecting the P4 mean curvature map projected onto the P32 brain (Fig. 2b). The P32 brain is significantly folded, which hides what happens inside sulci: a flattened P32 neocortex facilitates inspection across the entire neocortical surface (Fig. 2c). Although the P4 neocortex does not have regions of negative mean curvature, its curvature is heterogeneous. In particular, the hemispheric rim, close to the sagittal plane, is among the most curved regions. Projected into the P32, these regions of high curvature (red) become more often a gyrus, whereas those of lower curvature (blue) can become a sulcus or a gyrus. In addition the directions of curvature (indicated by arrows) appear to match the main axes of folding.

### The orientation of mature neocortical folding is aligned with the curvature directions of the unfolded neocortex

The orientation of folds at P32 was strongly aligned with the principal curvature directions of the unfolded P0 and P4 neocortices. In mature brains, the 2nd principal curvature direction runs along the main axis of each fold; the 1st principal direction cuts across folds (Fig. 3b). We quantified alignment as the cosinus of the angle between the 2nd principal curvature directions at corresponding points in the P0 and the P32 neocortices (Fig. 3a, 0=orthogonal; 1=perfect alignment). Alignment was substantial from birth and did not increase markedly with age, indicating a persistent geometric constraint on fold orientation (Fig. 3c, d. P0: 0.794±0.009; P4: 0.815±0.008; P8: 0.794±0.006; P16: 0.849±0.005; standard error by block jackknife; significance by spin tests; Figs. 3e-f).

**Figure 3.**
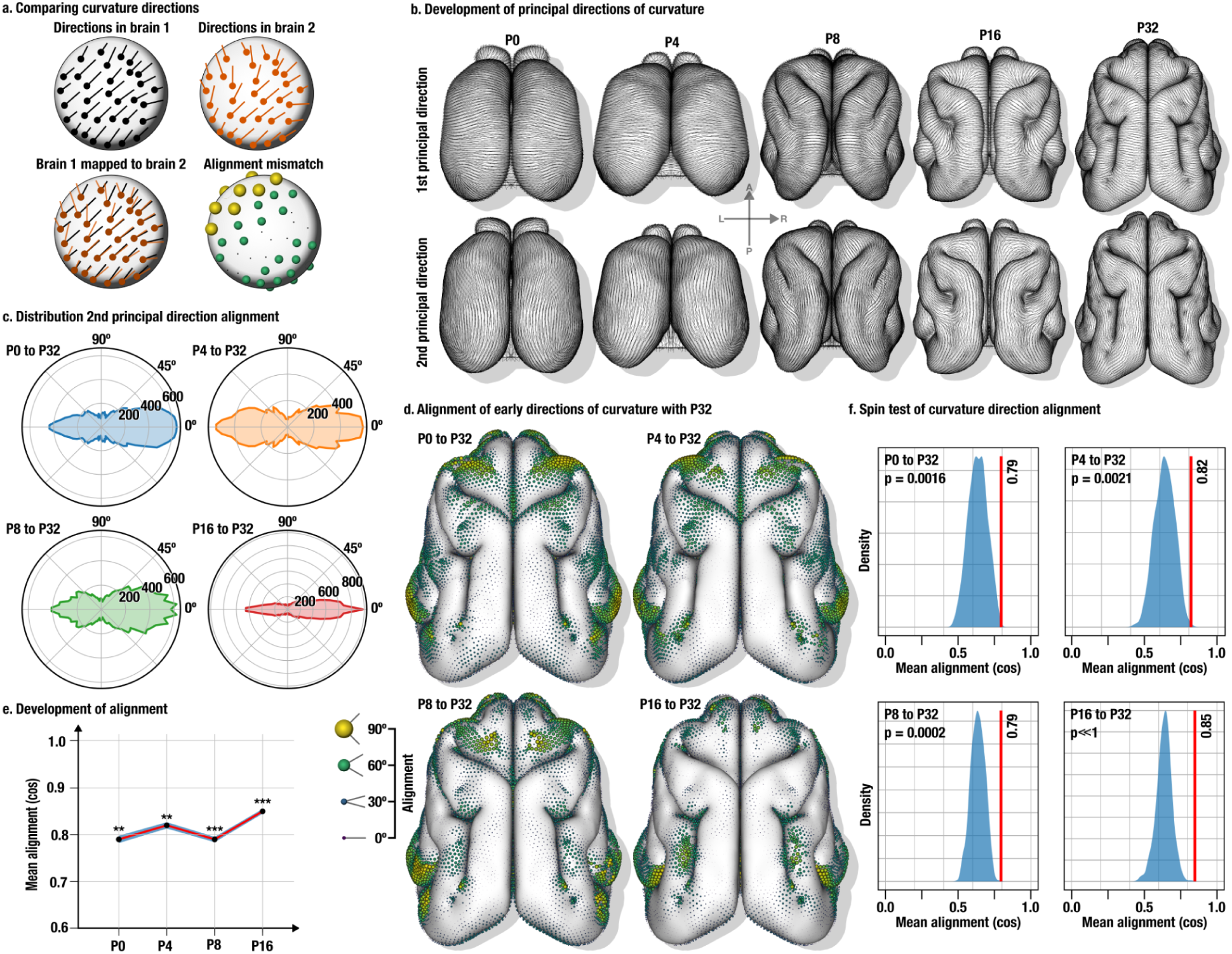
Alignment of folding orientation with pre-folding curvature directions. The orientation of mature folds, described by the 2nd principal curvature direction, is strongly aligned with the 2nd principal curvature direction of the unfolded newborn neocortex. **a.** Procedure for comparing curvature directions across developmental stages using surface registration. Directional discrepancy is visualised as sphere size at each vertex: small spheres indicate alignment; large spheres indicate misalignment. **b.** Maps of 1st and 2nd principal curvature directions from P0 to P32. The 2nd direction (bottom row) matches mature folding orientation with high accuracy throughout development. **c.** Alignment of the 2nd curvature direction between early stages and P32 is substantial from birth. **d.** Alignment maps comparing each stage with P32. **e.** Alignment values ± standard error (block jackknife). **f.** Significance estimated using spin tests.

### Generalisation across ages and individuals

The main findings generalised across individuals and across ages. We replicated all analyses in 21 additional ferret surface reconstructions (three individuals per age at P0, P2, P4, P8, P16, P32 and adult; Fig. 4a). Different deformation-minimising developmental trajectories were computed using random samples of subjects.

**Figure 4.**
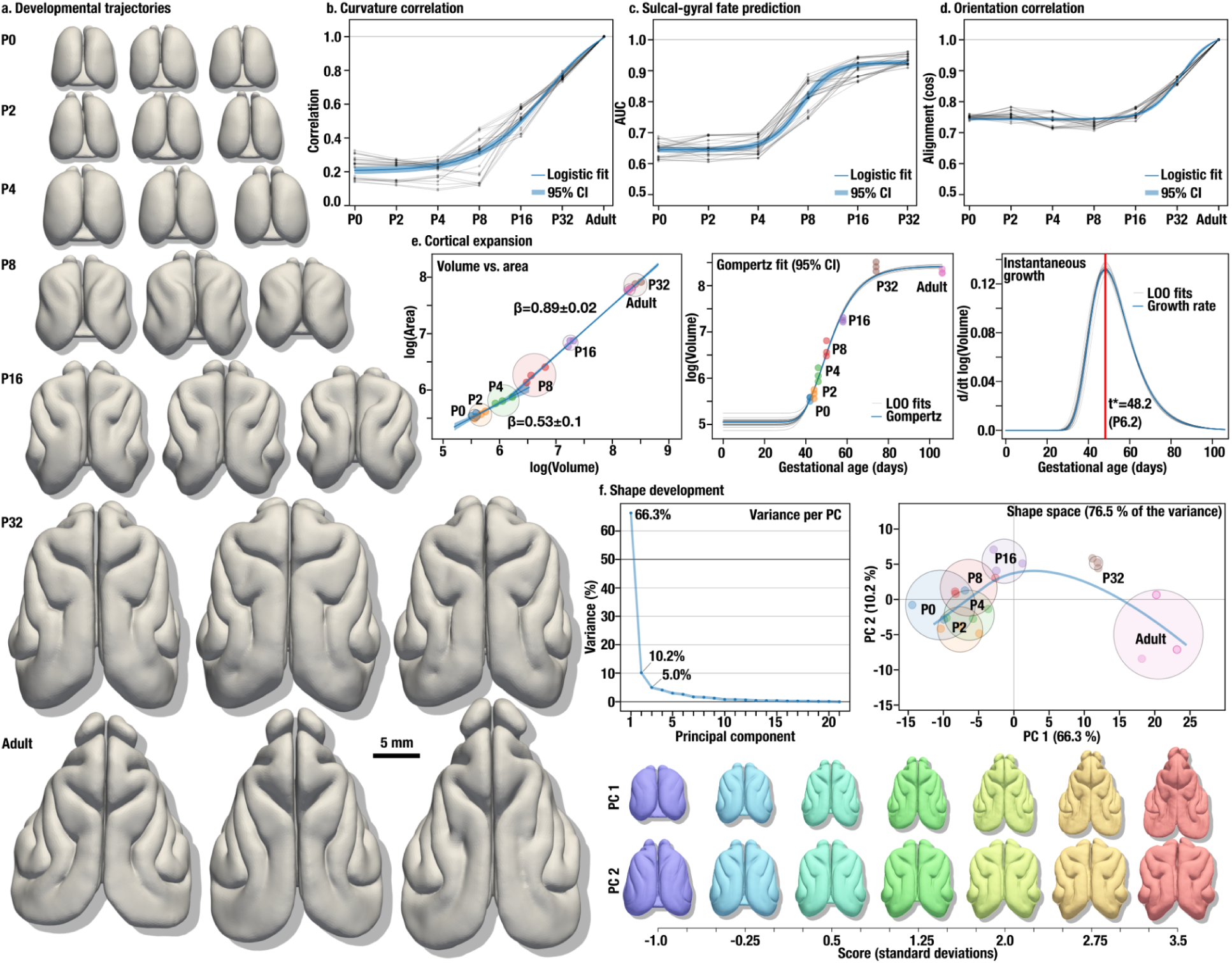
Generalisation across subjects. **a.** Extended sample of 21 ferrets spanning newborns to adults (3 individuals per age at P0, P2, P4, P8, P16, P32 and adult). **b.** Replication of the correlation between pre-folding and mature curvature. **c.** Replication of sulcal-gyral fate prediction, including the transition at the onset of folding. **d.** Replication of fold orientation alignment. **e.** Scaling of neocortical surface area with brain volume reveals two regimes: before folding, area scales as the square root of the volume (ß=0.53±0.10); after folding, the relationship approaches linearity (ß=0.89±0.02). A Gompertz growth fit to volumetric data estimates peak brain growth at approximately P6.2. **f.** Principal components analyses of volume-normalised surface reconstructions. The first 2 components capture 76.5% of variance. PC1 (66.3%) summarises the main developmental trajectory; PC2 (10.2%) captures variability in the extent of the frontal cortex, particularly around the cruciate sulcus. Circular zones contain subjects of the same age.

The two-phase pattern of curvature correlation was reproduced (Fig. 4b): a baseline of approximately r=0.2 before folding (P0, P2, P4), rising to substantially higher values after folding (P8, P16, P32). Sulcal-gyral fate prediction showed the same progression (Fig. 4c): AUC∼0.65 before folding, exceeding 0.8 after folding onset. Fold orientation alignment was consistently above 0.75 from the earliest stages (Fig. 4d).

Scaling of neocortical surface area with brain volume was markedly biphasic (Fig. 4e): a negative scaling regime before folding (ß=0.53±0.10) gave way to a near-linear regime after folding (ß=0.89±0.02). A Gompertz model fit to the volumetric data was consistent with the previously reported longitudinal data from Garcia et al. (2024) for later stages (P8 to adult). The estimated peak brain growth rate was at approximately P6.2, coinciding with the transition from the low-prediction to the high-prediction regime for sulcal-gyral fate.

Principal component analysis of volume-normalised surface reconstructions showed that the first two components capture 76.5% of shape variance (Fig. 4f; PC1: 66.3%; PC2: 10.2%). Despite volume normalisation, a residual effect of brain volume was apparent, likely reflecting non-linearity in the developmental trajectory. PC2 captured variability in frontal cortex extent, particularly around the cruciate sulcus.

### The curvature of the unfolded neocortex correlates with differential gene expression

We investigated the relationship between pre-folding curvature, early gene expression, and sulcal-gyral fate by reanalysing data from De Juan Romero et al. (2015) and Singh et al. (2024). De Juan Romero et al. (2015) showed that gene expression at P0 and P2 in ferret correlates with the later positions of gyri and sulci at P6, supporting the hypothesis of a genetically encoded protomap (Rakic 1988). Singh et al. (2024) extended this analysis to embryonic stages (E30 and E34, approximately 2 weeks before birth at E42), demonstrating significant molecular differences between regions destined to become the lateral sulcus (LS) and the splenial gyrus (SG).

Pre-folding curvature and gene expression were significantly correlated. We analysed 9 pre-folding ferret surface reconstructions (3 at P0, 3 at P2 and 3 at P4; Fig. 5a), computing mean curvature profiles in parasagittal slices (Fig. 5d) comparable to those of De Juan Romero et al. (2015). In-situ hybridisation data were available for *Fgfr3* at P0, *Cdk6* at P2, and *Eomes* at P6 (the last also providing the sulcal-gyral reference; Fig. 5c, e). Mean curvature profiles at P0 correlated with *Cdk6* expression at P2 (r=0.31, p<0.05) and *Eomes* expression at P6 (r=0.57, p<0.01). As in our previous analyses, the prediction of sulcal-gyral fate at P6 from curvature at P0 was statistically significant (AUC=0.66). Conversely, expression of *Fgfr3* at P0 correlated with curvature at P2 (r=0.66, p<0.01) and P4 (r=0.28, p<0.01). The prediction of sulcal-gyral fate from gene expression patterns, however, was not statistically significant. Mediation analysis indicated that the direct effect of curvature at P0 was the main determinant of sulcal-gyral fate at P6 (Model A, Fig. 5g; curvature odds ratio = 2.49). Gene expression at P2 contributed only marginally (odds ratio = 0.97).

**Figure 5.**
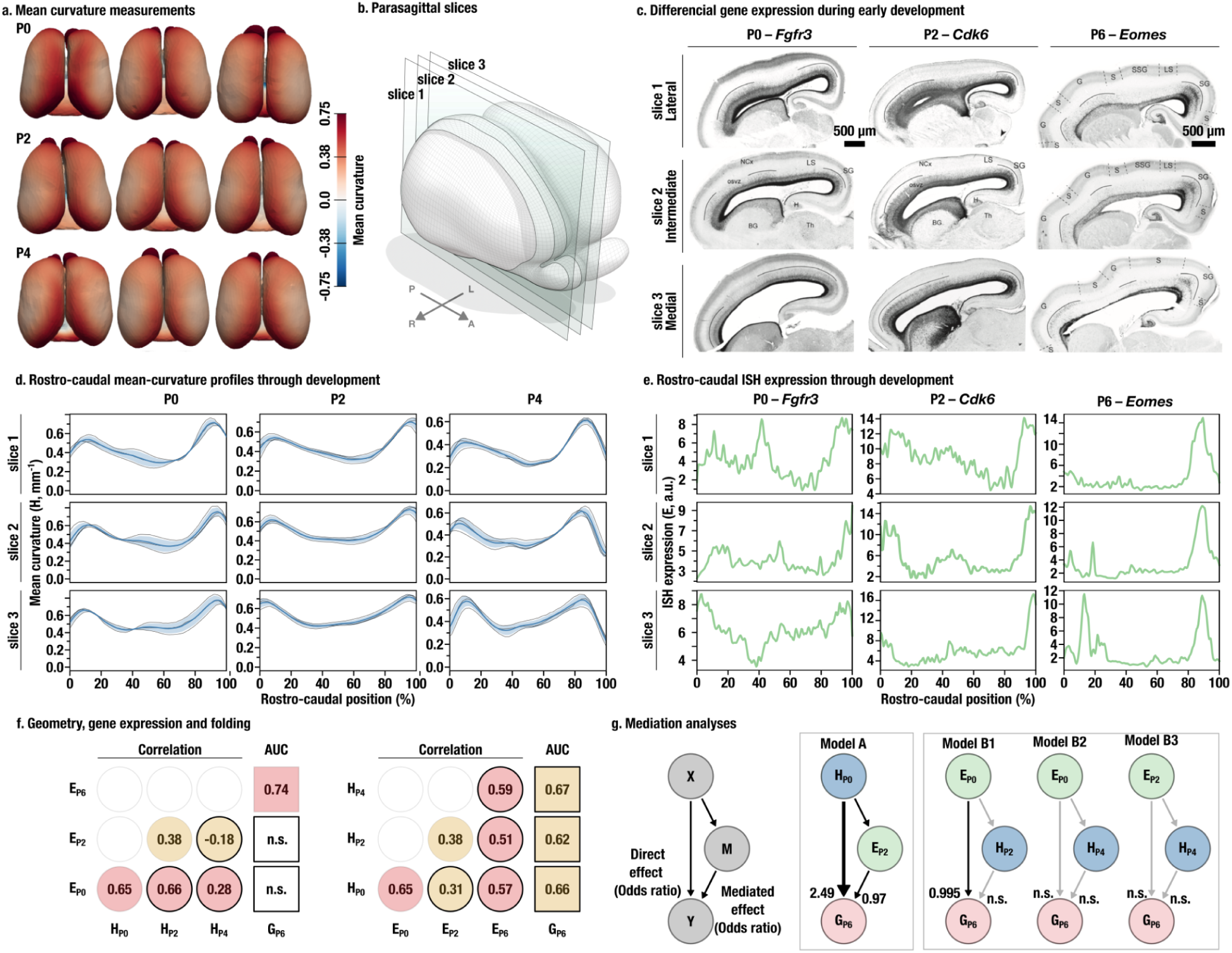
Pre-folding geometry is a stronger predictor of sulcal-gyral fate than early differential gene expression. **a.** Mean curvature measurements on pre-folding surface reconstructions. **b.** Parasagittal slices used for mean curvature profiles and gene expression analysis, comparable to those in De Juan Romero et al. (2015). **c.** Maps of differential gene expression at P0, P2 and P6 from De Juan Romero et al. (2015). **d.** Rostro-caudal profiles of mean curvature through development. **e.** Rostro-caudal profiles of gene expression through development. **f.** Correlation between curvature profiles, expression profiles, and sulcal-gyral fate at P6. Circles without border: same time; with border: different time; squares: prediction. Background orange: p<0.05, red: p<0.01. **g.** Mediation analysis. Model A: curvature at P0 (H_P0_) predicts sulcal-gyral fate at P6 (G_P6_) with mediation through expression at P2 (E_P2_); direct effect odds ratio = 2.49. Model B: expression as primary predictor, mediated by later curvature; only model B1 yields a weak statistically significant direct effect of P0 expression (odds ratio=0.995).

Notably, both the curvature-fate and expression-fate correlations were driven principally by signal in the presumptive lateral sulcus and splenial gyrus. The position of the splenial gyrus corresponds with the high curvature region pre-folding, and the position of the lateral sulcus corresponds with the low curvature region pre-folding (Fig. 2b-d). This low curvature region will also be the place where the supra-sylvian gyrus (SSG) develops. De Juan Romero et al. (2015) reported that gene expression peaks capture the splenial gyrus but not the SSG. The work of Singh et al. (2024) does not discuss this gyrus and only focuses on the difference between LS and SG.

The global shape of the embryonic ferret brain is similar to that of a P0, and the regions of high and low curvature are largely comparable. Inspection of published 2D images from Singh et al. (2024) revealed a significant difference in curvature of the ventricular zone and pial contours between the presumptive lateral sulcus (low curvature) and the splenial gyrus (high curvature), consistent with our postnatal findings (Mann-Whitney U = 0, rank-biserial r = −0.87, p≪1). Two-dimensional curvature measurements from the images of De Juan Romero et al. (2015) yielded results comparable to those from 3D reconstructions.

### A mechanical model predicts the role of early high-curvature regions in organising the brain folding

The initial geometry of the ferret neocortex has higher curvature along the hemispheric rim than on the lateral surface. If the curvature of the cortex were the same everywhere, and if cortical growth is sufficient to make the cortex buckle and fold, the position of these folds should be random. However, if there are regions with high and low curvature, those with lower curvature should fold first, priming the dynamics of the folding process and canalising the folding pattern. We built a series of 2D mechanical models to test whether this curvature heterogeneity is sufficient to organise the folding pattern. We compared two geometries: cortices with uniform curvature (circular models, Fig. 6a) and cortices with high-curvature flanking regions of medium and low curvature (trikyklon models, from Greek *tri:* three and *kyklos:* circle, Fig. 6b). Cortical thickness was constant within each model and varied within the same range across models. Material properties, total growth, and outer perimeter were held constant. The cortex and the core were modelled as hyper-elastic materials (Goriely 2017).

**Figure 6.**
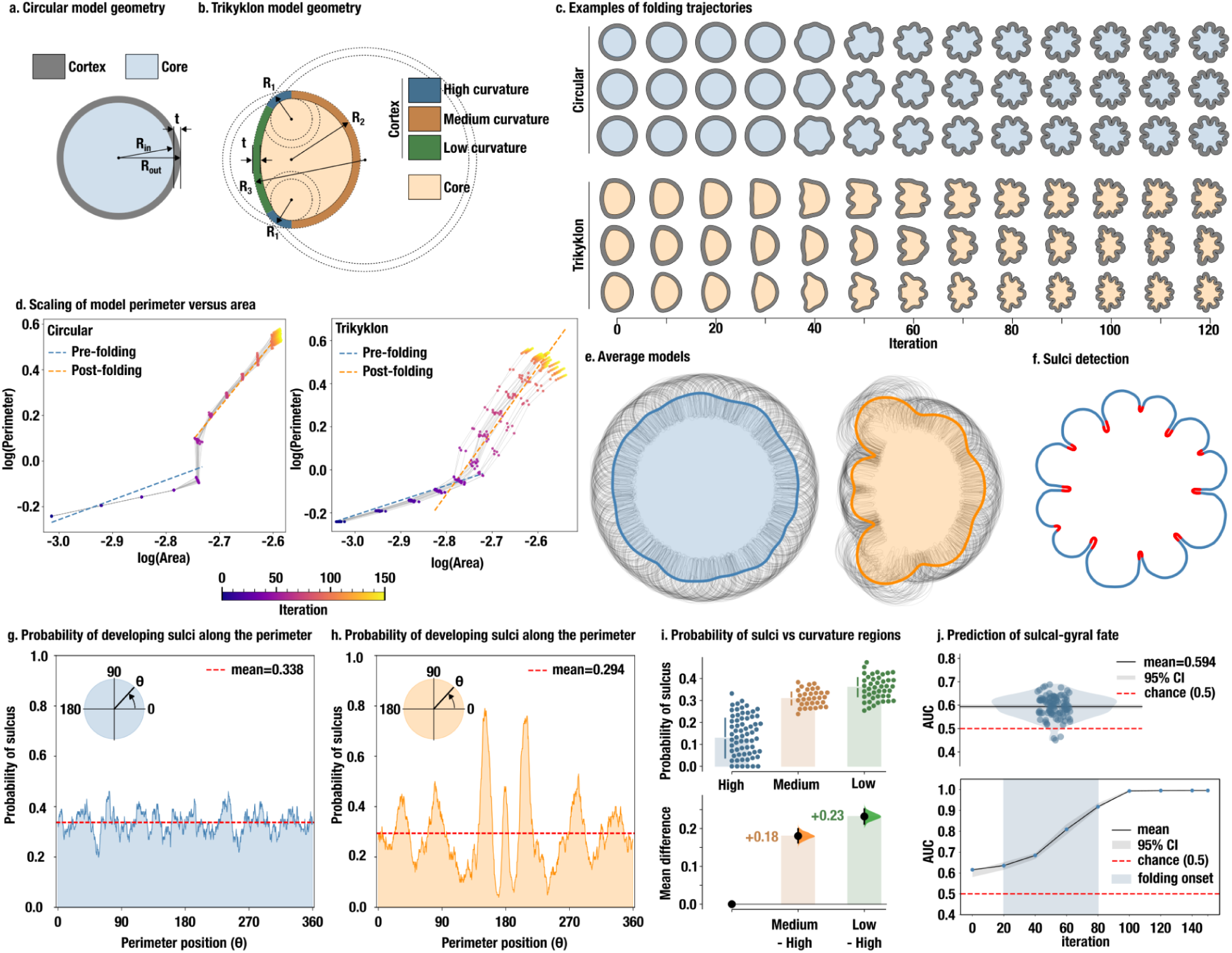
A mechanical model demonstrates how regions of high-curvature guide the emerging folding pattern. **a.** Circular reference geometry (uniform curvature). **b.** Trikyklon geometry: 3 distinct curvature regions assembled from circles of different radii (2 small circles with high curvature, one medium, and one large with low curvature). Cortical thickness is constant within each model. **c.** Three representative simulations for circular (top) and trikyklon (bottom) geometries. **d.** Perimeter versus area scaling shows two phases – pre-folding and post-folding – in both the simulations and the ferret data. **e.** Averaging 100 circular models produces a nearly circular shape, indicating that fold positions are random under uniform curvature. Averaging 100 trikyklon models reveals a consistent pattern, with high-curvature regions remaining gyral and organising folds in adjacent regions. **f.** Automated labelling of sulcal regions (sulci: red, gyri: blue). **g.** Sulcal probability is uniform along circular model contours. **h.** Sulcal probability is markedly non-uniform in trikyklon models: high-curvature regions strongly resist sulcal fate. **i.** Probability of sulcal fate in high-, medium-, and low-curvature regions across simulations. **j.** Prediction of final sulcal-gyral status from initial curvature: distribution of AUC (top), trajectory of AUC (bottom).

All models grew without folding initially. After a critical growth threshold was reached, the cortex buckled, sometimes even twice (Fig. 6c). Thinner models folded earlier and more extensively, as expected in theory (Toro and Burnod 2005, Bayly et al. 2013) and matching observations from cross species folding (Heuer et al. 2019, Heuer et al. 2023). Perimeter versus its area plots showed a biphasic pattern mirroring that of the ferret data (Fig. 6d). The average of 100 circular models was nearly circular (Fig. 6e, left): fold positions were random. The average of 100 trikyklon models showed a consistent, non-random pattern: high-curvature regions remained gyral and acted as anchors, framing and organising the surrounding folds (Fig. 6e, right). We labelled sulci automatically based on the mean curvature of the contours (Fig. 6f). Sulcal probability was uniform along circular model perimeters (Fig. 6g) but markedly patterned in the trikyklon models (Fig. 6h–i): high-curvature regions were significantly less likely to develop sulci, with the exception of models with very thin cortices.

As in the ferret brains, the curvature of the initial models predicted the sulcal-gyral state of the final model (Fig. 6j top). The AUC is moderate but highly significant (mean AUC=0.594, 95% CI [0.585, 0.603], p≪1) with only a few models having an AUC below chance level. This is because high-curvature regions only guide the final folding pattern but do not determine it completely: the final folding pattern appears after buckling, at which point the prediction accuracy jumps to become very high (Fig. 6j bottom).

## Discussion

The neocortex is the phylogenetically most recent structure of the mammalian brain and underlies the most distinctive human cognitive functions, including language, symbolic communication, and abstract reasoning (Jablonka and Lamb 2024, Namba and Huttner 2024). Understanding the nature and origin of its organisation remains a central challenge in neuroscience, and a growing body of work highlights an important role played by folding (Del Valle Anton and Borrell 2022). Work over the last three decades has demonstrated the fundamental role of early molecular gradients in arealisation (Mallamaci et at 2000, Fukuchi-Shimogory and Grove 2001, Caronia-Brown et al. 2014) and in the establishment of a molecular gyral protomap specifying the position of future folds (De Juan Romero et al. 2015, Singh et al. 2024). Our analyses show that the geometry of the unfolded neocortex provides a complementary signal with comparable, and by some measures greater, predictive power. The principal curvature directions of the unfolded newborn neocortex constrain the orientation of mature folds from the first day of postnatal life; early regions of high curvature act as a scaffold over which sulci and gyri organise; and curvature at P0 correlates with the patterns of differential gene expression at P2 and P6. This finding raises the possibility that early geometry may partly regulate differential gene expression rather than merely accompanying it.

The strong geometric constraint on fold orientation deserves particular attention. Unlike the prediction of sulcal-gyral fate, which strengthens progressively as folding begins, fold orientation alignment is substantial and essentially constant from birth. Fold orientation determines the trajectory of short-range associative fibres (U-fibres) that connect gyral crowns (Yoshino et al. 2020, García et al. 2021). If orientation is set by early brain geometry before connectivity is established, the architecture of local cortico-cortical circuits may be, in part, a geometric consequence rather than a primary molecular specification.

### Mechanical interpretation

Our model provides a parsimonious explanation for these observations. The neocortex folds due to the larger growth of the grey matter relative to the white matter; beyond a critical threshold, this configuration is mechanically unstable and the cortex buckles (Toro and Burnod 2005; Tallinen et al. 2014, 2016; Greiner et al. 2021). The position and orientation of individual folds are regulated by the initial geometry: regions of lower curvature require less growth to reach the buckling threshold and therefore fold first; regions of higher curvature resist folding and act as anchors. This is because curvature delays the wrinkling instability: bending stiffness is effectively increased by curvature, so greater growth is needed to initiate folding on a curved surface than on a flat one (Budday et al. 2015; Jia and Goriely 2018). This geometric mechanism was first proposed by Todd (1982) and explored *in silico* by Toro (2012) and Tallinen et al. (2014, 2016). Our trikyklon simulations demonstrate explicitly that even modest curvature heterogeneity is sufficient to impose a reproducible, non-random folding pattern.

In both our models and in ferrets, early curvature provides only a moderate prediction of the final folding pattern (average AUC∼0.6 in models; AUC∼0.65 in ferrets). This reflects the fact that the complete folding configuration emerges only after buckling, with early geometry serving primarily as a guide. We therefore expect the predictive power of early brain geometry to be higher in species with relatively simple, less folded brains – a pattern consistent with our simulations. This interpretation contrasts with gyral protomap hypotheses, which posit that differential gene expression patterns at early stages should already specify the full mature folding pattern.

The large-scale geometric reorganisation produced by folding may then generate further positional signals relevant to subsequent developmental events, including the organisation of connectivity (García et al. 2021; Heuer and Toro 2019; Heuer et al. 2025) and neural activity patterns (Pang et al. 2023; Schwartz et al. 2023).

### Geometry and mechanics within the broader field

The question of what determines neocortical organisation has most often been framed as a trade-off between intrinsic genetic programmes and extrinsic experience-dependent processes, with thalamo-cortical afferents as the principal extrinsic regulator (Krubitzer 2007, Vue et al. 2013, Chou et al. 2013). Geometry and mechanics occupy an interesting intermediate position. Like genetic programmes, they are intrinsic to the developing brain; like experience-dependent processes, the physical laws governing mechanical instabilities are common to all individuals regardless of their specific genetic backgrounds. The non-linearity of developmental mechanics means that nominally homogeneous growth can produce complex, reproducible structures in a manner that is decoupled from any particular genetic instruction (Foubet et al. 2019, Heuer and Toro 2019). This suggests that mechanical morphogenesis may constitute a third fundamental axis of neocortical organisation – alongside genetics and experience-dependent processes – as recently proposed from comparative neuroanatomical evidence (Heuer et al. 2025).

### Interaction between molecular positional information and self-organisation

Morphogen gradients providing positional information are well established as central regulators of embryonic patterning (Tkačik and Gregor 2021) and neocortical arealisation (Mallamaci et at 2000, Fukuchi-Shimogory and Grove 2001, Caronia-Brown et al. 2014, De Juan Romero et al. 2015, Singh et al. 2024). Self-organisation through emergent reaction-diffusion mechanisms provides a complementary class of patterning (Turing 1952), and recent work suggests that these two processes interact to distribute patterning across spatial scales (Tzika et al. 2023). Mechanical morphogenesis and global curvature occupy a similar intermediate position: they can arise from either top-down or bottom-up processes, but once established, they integrate local variations into a global constraint essentially instantaneously. The propagation of mechanical deformation across the whole brain is orders of magnitude faster than the diffusion of signalling molecules, making it an efficient medium for large-scale integration. In the model we propose, local differential growth triggers a global mechanical instability that produces a sudden, non-linear shape change: the folding event itself. The wavelength of individual folds is set by local tissue properties and growth dynamics, but their global arrangement is constrained by the entire neocortical geometry. Mechanical forces may therefore mediate a continuous bidirectional dialogue between gene expression, morphology, and the subsequent regulation of gene expression.

### Limitations

Several limitations should be kept in mind. Our dataset is cross-sectional: we compare different individuals at different ages rather than tracking single subjects through development. Longitudinal imaging by MRI or functional ultrasound would allow within-individual tests of our hypotheses, but acquiring such data in newborn ferret pups involves a trade-off between resolution and the difficulty of anaesthetising neonates. The stability of the ferret folding pattern across individuals makes cross-sectional comparisons informative, but longitudinal studies would strengthen causal inference.

Second, the gene expression data were not acquired from the same individuals used for MRI, and the embryonic curvature estimates rely entirely on published 2D images. The conclusions of the mediation analysis should therefore be regarded as preliminary. The remarkable reproducibility of molecular patterns across ferret individuals (De Juan Romero et al. 2015; Singh et al. 2024) gives some confidence that the broad findings will generalise, but verification using 3D embryonic reconstructions integrated with spatially registered gene expression data is an important next step. Finally, a perturbation experiment, as proposed in Foubet et al. (2019) and Heuer and Toro (2019), should provide more direct proof of causality.

Third, a complete characterisation of the role of early geometry would require histological data on cortical thickness and cell density, measurements of tissue mechanical properties, and information about the development of connectivity – none of which are currently available in our analysis. A given surface shape can be associated with very different internal stress distributions, and geometric curvature alone does not fully characterise the mechanical state of the tissue (Xu et al. 2010). Acquiring and integrating these modalities remains a substantial experimental and computational challenge.

### New perspectives

Several questions emerge from our findings. The cortical hem, antihem, and anterior neural ridge – major signalling centres that release morphogens such as Wnts, BMPs, FGFs, and SHH – are positioned at points of high curvature along the edges of the early neocortex (Subramanian et al. 2009, Caronia-Brown et al. 2014). These secreted cues regulate graded expression of transcription factors including Emx2 and Pax6. Whether the locations of these centers are determined by, or help produce, local curvature remains unresolved. If morphogen gradients correlate with neocortical shape, molecular and geometric positional information may not be independent but mutually reinforcing – a possibility that could be tested by combining morphogen imaging with curvature mapping in the early embryo.

The early geometry we describe may itself reflect consequences of much earlier processes. The shape established during neural tube folding and telencephalic expansion is mechanically regulated, and could be the distal cause of the curvature patterns we observe postnatally. Under this interpretation, the positional encoding of folding patterns begins earlier in development than current molecular protomap models assume.

The broad similarity of early brain geometry across gyrencephalic and lissencephalic species is informative. Species that do not develop folds may simply not grow large enough for cortical expansion to exceed the buckling threshold. Primary gyri form once this threshold is exceeded; secondary gyri form when the crowns of primary gyri exceed the threshold in turn, generating a hierarchy of geometric constraints on neocortical organisation. An analogous argument may apply to other folded cortices, including the cerebellum and hippocampus.

### Testing and falsification

The supra-sylvian gyrus (SSG) provides an instructive test case. The SSG occupies a region of intermediate curvature in the pre-folding brain and lacks a specific molecular signature in existing protomap studies (De Juan Romero et al. 2015). Our geometric model predicts that the SSG forms without prior molecular specification, driven by the folding dynamics of surrounding gyri and sulci. A demonstration of an early (for example, P0) molecular protomap domain for the SSG would challenge this prediction. More broadly, such a finding would raise the question of whether the molecular wavelength of protomap patterns and the mechanical wavelength of the folding instability evolved to match, producing a system in which both encodings cooperatively specify the same pattern.

### Relation to earlier work

Our results call for a reassessment of the role of molecular signals as the sole determinants of the folding protomap. The geometric hypothesis does not dismiss the work of De Juan Romero et al. (2015) and Singh et al. (2024), which reveals real and reproducible molecular differences between prospective gyri and sulci. Rather, our data suggest that these molecular correlates of folding may partly reflect a cellular response to early geometric signals rather than their primary cause. This reinterpretation places progressively more demanding requirements on molecular protomap models as brain complexity increases: in macaques or humans, whose neocortices carry dozens of consistently placed folds, an increasingly precise and spatially complex molecular specification would be needed. The possibility that early geometric constraints reduce this molecular specification burden – by providing the primary positional scaffold – is worth examining in more gyrencephalic species.

## Conclusion

We have shown that the early geometry of the neocortex encodes substantial information about the position and orientation of adult folds before any folding has begun. Pre-folding curvature predicts sulcal-gyral fate and fold orientation with statistical significance, correlates with early patterns of gene expression, and is identified by mechanical simulation as the principal organisational cue for the emerging folding pattern. These findings suggest that neocortical geometry constitutes a source of positional information that acts alongside molecular gradients in encoding the adult cortical architecture. Mechanical studies alone, or molecular studies alone, provide an incomplete picture; an adequate understanding of cortical development and evolution will require the integrated characterisation of molecular, geometric, and mechanical processes and their dynamic interplay.

## Methods

### MRI data collection and surface reconstruction

We collected ex-vivo MRI data from developing ferret brains and generated high-quality, manually edited cortical surface reconstructions. A cross-sectional developmental series was assembled at P0, P2, P4, P8, P16, P32, and adult to reconstruct developmental trajectories from birth to adulthood. These trajectories provided point-to-point correspondences between the P0 and adult neocortices.

High-resolution MRI were acquired *ex vivo* using a small animal 7 Tesla Bruker MRI scanner (Neurospin, Saclay, France). The acquisitions were performed postmortem in order to improve sensitivity. Ferrets were euthanised by an overdose of pentobarbital and perfused transcardially with 0.9% saline solution and post-fixed with phosphate-buffered 4% paraformaldehyde (PFA). After extraction, brains were stored at 4°C in a 4% PFA solution until the MRI acquisition. All procedures were approved by the Institutional Animal Care and Use Committee of the Universidad Miguel Hernández and CSIC (Consejo Superior de Investigaciones Científicas), Alicante, Spain.

Brains were transferred to a 0.01 M phosphate-buffered saline (PBS) solution for rehydration 24 hr before MRI acquisition. Shortly before MRI acquisition, the brain sample was transferred to a plastic tube filled with nonprotonic liquid (fluorinert) in order to avoid air-tissue interfaces that may induce susceptibility artefacts, as well as to avoid foldover MRI artefacts due to a proton signal coming from a protonic liquid outside the imaging field of view (McRobbie, Moore, & Graves, 2017). The tube was then placed in a dedicated holder in the middle of the transmit/receive MRI volume radiofrequency coil. Temperature stability was ensured by a regulated room temperature as well as the cooling of the gradient coils, by water at 16°C that was constantly flowing inside the innermost part of the magnet. The equilibrium temperature at the sample was 20°C. High-resolution T2-weighted MRI data were acquired using a multi-slice multi-echo (MSME) sequence with 18 echo times and 0.12 mm isotropic voxels.

### Surface reconstruction and neuroanatomical measurements

We used several tools to produce cortical surface reconstructions from the MRI data. We reoriented the data to align the brains with respect to the stereotaxic axes using Reorient (https://neuroanatomy.github.io/ reorient, Heuer et al. 2020). A first mask was obtained interactively using Thresholdmann (https://neuroanatomy.github.io/thresholdmann, Heuer et al. 2024), which allows the creation of space-varying segmentation thresholds. These masks were then refined using BrainBox (https://brainbox.pasteur.fr, Heuer et al. 2016). BrainBox is a Web application providing several tools for collaborative manual segmentation of MRI data. The main task in BrainBox was to make sure that sulcal regions were open, and to correct additional segmentation errors. The masks generated by BrainBox were then used to produce surface meshes, using the implementation of the marching cubes algorithm provided by Scikit-Image (van der Walt, 2014). The meshes were then regularised using Graphite (https://github.com/BrunoLevy/ GraphiteThree), which also automatically corrects a certain number of topological errors. Finally, the topology of the meshes was manually edited to obtain meshes of genus 0 (topologically spherical) using MeshSurgery (https://github.com/neuroanatomy/MeshSurgery.

A series of neuroanatomical measurements were obtained from these meshes, including brain volume, cerebral volume, surface area, and gyrification index, using the command line tool meshgeometry (https://github.com/neuroanatomy/meshgeometry).

### Curvature computation

Cortical surfaces at postnatal ages P0, P2, P4, P8, P16, P32, and adult were represented as triangulated meshes reconstructed from MRI data. Mean curvature H=(k_1_+k_2_)/2 and Gaussian curvature G=k_1_k_2_ were estimated at each vertex from the two principal curvatures k_1_, k_2_ computed by fitting a local quadric over a geodesic neighbourhood. Curvature values were clipped to ±5 standard deviations from the median and subsequently smoothed by iterative graph-Laplacian diffusion to suppress mesh noise.

### Surface Registration

For a quantitative comparison across individuals we used a non-linear surface matching algorithm which builds a mapping from one native surface to the other using a reduced number of eigenvectors from the source mesh Laplacian. A linear combination of these eigenvectors was found which minimised metric distortions while maintaining smoothness (see Heuer et al. 2025 for more details). To compare features across ages, scalar and vector fields were transferred between meshes using precomputed barycentric mappings derived from sphere-based inter-individual registration. Each early-age field was resampled onto the adult whole-brain topology, then projected onto a regularised hemisphere mesh. A manually delineated neocortex mask, transferred across adult individuals by the same sphere-based resampling, was applied throughout: vertices outside the neocortex were excluded from all statistical analyses.

### Spin Tests

Statistical significance of spatial correspondences was assessed using spin tests, a form of spatial permutation test that preserves the autocorrelation structure of cortical maps (Alexander-Bloch et al. 2018, Markello et al. 2022). For each of 1000 permutations, a uniform random rotation was applied to the spherical parameterisation of the adult hemisphere, permuting the vertex ordering of the adult reference map while leaving its spatial structure intact. The observed statistic was compared to the resulting null distribution and a p-value computed as the proportion of null statistics exceeding the observed value. A kernel density estimate of the null distribution was also used to obtain an interpolated p-value. Standard errors were estimated by block jackknife over 100 spatial blocks to account for residual autocorrelation.

### Curvature correlation test

To test whether the curvature pattern at each early age predicts the adult folding pattern, we computed the Pearson correlation between the early-age mean curvature map (resampled to adult topology) and the adult mean curvature map, restricted to neocortical vertices. Statistical significance was assessed by a spin test of the adult reference map (1000 permutations). For the cohort analysis, this procedure was repeated independently for each of 21 ferret developmental trajectories and both hemispheres (left and right), and individual effect sizes were combined in a meta-analysis (see below).

### Sulco-gyral fate prediction

To test whether early curvature predicts whether a cortical region will become a sulcus or a gyrus in the adult brain, we trained a logistic regression classifier using two curvature features – early-age mean curvature H and Gaussian curvature G – to predict the binary adult folding identity (sulcus: adult mean curvature<0; gyrus: adult mean curvature>0). The regularisation parameter was selected by 5-fold cross-validated AUC maximisation. Predictive accuracy was quantified by the area under the ROC curve (AUC); a bootstrap procedure (1000 resamples) provided a standard error for the AUC. Statistical significance was assessed by a spin test: for each of 1000 permutations, the adult sulcus/gyrus labels were rotated and the classifier was retrained with fixed regularisation, providing a null AUC distribution. For the cohort analysis, this was repeated across all trajectories and hemispheres and combined in a meta-analysis.

### Folding direction alignment test

To test whether the direction of early folding predicts the adult folding orientation, we compared the second principal curvature direction field d_2_ at each age against the adult reference field. Direction fields were transferred to a common spherical reference frame: each tangent-plane direction vector was mapped to a displacement on the unit sphere via barycentric interpolation of the spherical parameterisation. Because principal curvature directions are defined up to sign, alignment was measured as the mean absolute cosine similarity, A = < |s^(0)^ • s^(1)^|>, over neocortical vertices (A = 0 for orthogonal, A = 1 for perfect alignment). Statistical significance was assessed by a spin test of the early-age direction field on the sphere (1000 uniform random rotations). For the cohort analysis, individual trajectory results were combined in a meta-analysis.

### Cohort meta-analysis

Effect sizes ($\rho$ for curvature correlation, AUC for sulco-gyral fate, mean absolute cosine A for orientation) and their standard errors were computed independently for each trajectory and hemisphere. Results were combined using inverse-variance weighted meta-analysis. A fixed-effects estimate was computed as the weighted mean θ_FE_=∑w_i_ θ_i_ / ∑w_i_ (with w_i_ = 1/σ_i_^2^). Between-study heterogeneity was quantified by Cochran’s Q and the I^2^ statistic. A random-effects estimate was obtained by the DerSimonian–Laird moment estimator, which adds the between-study variance 𝜏^2^ to the within-study variance before computing weights. Both fixed- and random-effects p-values were computed from z-scores against the standard normal distribution.

### Alignment to published ISH Contours

Three sagittal cross-sections were extracted per specimen (lateral, intermediate, and medial slices). The longest contiguous neocortex-labelled arc of each cross-sectional contour was retained and parametrised by normalised arc length. In-situ hybridisation (ISH) gene expression data were taken from De Juan Romero et al. (2015, Fig. 6), which reports *Fgfr3* expression at P0, *Cdk6* at P2, and *Eomes* at P6. Pial and SVZ contours were digitised from the figure’s SVG annotation file and resampled to N = 200 equally-spaced arc-length points. Each mesh neocortex arc was aligned to the corresponding published pial contour via a linear arc-length mapping, x_djr_ = m t_mesh_ + n, with parameters (m, n) determined per specimen per slice by visual inspection (36 parameter pairs total).

### Curvature and expression profiles

Mean curvature H_s_ was interpolated onto each slice contour by linear edge interpolation and resampled to the 200-point x_djr_ grid. Expression profiles were obtained by sampling red-channel pixel intensity from the ISH image along the digitised SVZ contour, inverting (E = 255 - I_raw_), and applying three iterations of mild Gaussian smoothing. A single expression profile was shared among the three specimens of each age. Per-age mean and inter-specimen standard deviation were then computed across the three curvature profiles.

For each specimen and slice we computed: (i) Pearson correlation r(H, E) between curvature and expression; (ii) robust linear regression (Huber M-estimator) of curvature on expression; (iii) binned mutual information (15 bins, Miller–Madow bias correction). Per-age means and standard deviations across three specimens are reported.

Cross-age predictability was assessed by correlating each P0 specimen’s curvature profile with the P2 and P6 expression profiles (r(H_P0_, E_P2_) and r(H_P0_, E_P6_)). Formal mediation of the P0 curvature → later expression → sulcal fate pathway was tested using parametric bootstrap mediation (500 replications), with binomial GLM as the outcome model and OLS as the mediator model using the `statsmodels` package. Sulcal positions were defined by the intersection of radial pial–SVZ line segments with blue annotation marks in the published figure.

### Comparison of embryonic curvature

We digitised the boundaries of the Lateral Sulcus (LS) and Splenial Gyrus (SG) along the pial surface and ventricular zone (VZ) from two embryonic ferret brain sections (E30, E34) published in Singh et al. (2024). Paths were uniformly resampled using arc-length interpolation. To each resampled path we fitted a parametric quadratic B-spline and computed mean absolute curvature along the spline as κ=<|x’y”− y’x”| / (x’^2^+y’^2^)^3/2^>. Robustness was assessed by a bootstrap procedure: 1,000 iterations each using a random two-thirds subsample of the resampled vertices (drawn without replacement, spatial order preserved). LS versus SG were compared using two-sided Mann-Whitney U tests on the resulting bootstrap distributions; effect size was reported as the rank-biserial correlation r=Z/N^0.5^.

### Folding model

We used a 2D Material Point Method (MPM) to simulate the growth-driven folding of an elastic cortical shell. The cortex was modelled as a thin annular layer (Young’s modulus E_c_ = 3,000, Poisson’s ratio 𝜈 = 0.3) surrounding a softer core (E_k_ = 1,000, 𝜈 = 0.3). Growth was applied exclusively to the cortex, in the tangential direction only, with a time constant of 𝜏_tt_ = 150 ms, leaving the radial dimension unconstrained. Each simulation contained 300,000 MPM particles (150,000 cortex, 150,000 core) and ran for 150 growth iterations.

### Geometry

We compared two types of initial geometry, both with the same outer perimeter $P$. (1) Circular models: A ring of uniform curvature, with fixed outer radius R_out_, cortex thickness 𝛿 = 0.05. (2) Trikyklon models: A four-arc closed curve with two regions of high curvature (small-radius arcs, radius s) and one region of low curvature (large-radius arc, radius b). Five parameters (b, r, s, d, t) define the shape. For each of the 100 trikyklon simulations, parameters were drawn uniformly at random: b ∈ [6.0, 7.0], r ∈ [4.0, 5.0], s ∈ [2.0, 2.5], d = 3.5 (fixed), t ∈ [0.8, 1.2]. Each shape was rescaled to match P, ensuring that differences in folding patterns reflect curvature inhomogeneity rather than total tissue area.

### Batch simulations

For each geometry type, 100 independent simulations were run. Interior particle positions were generated by `numpy` rejection sampling with a per-simulation random seed, bypassing a limitation of the GPU random-number generator that would otherwise produce identical particle configurations across runs. To cancel biases introduced by the MPM grid’s discrete symmetry, each simulation was additionally rotated by a uniformly random angle before the simulation and counter-rotated after, so that all final contours are expressed in a common reference frame.

### Sulcus detection

In each simulation the outer contour was sampled at N = 1,000 points. A contour point was classified as lying in a sulcus if the locally smoothed signed curvature was positive (inward-facing). Specifically, at point i, the vectors a_i_ = p_i+D_ - p_i_ and b_i_ = p_i-D_ - p_i_ (with D = 10, indices mod N) were normalized and their 2D cross product was computed. The resulting signal was smoothed by 10 iterations of a circular box average. Points with positive smoothed curvature were labelled as sulci.

### Sulcus frequency maps

For each of the two batches, we computed the sulcus frequency at every contour point: the fraction of the 100 simulations in which that point fell inside a sulcus. In the circular batch this frequency should be spatially uniform (no preferred location); in the trikyklon batch we expected frequency to vary with the initial curvature of the shape. The initial curvature profile of the trikyklon contour was computed with the same signed-curvature formula and averaged across all 100 parameter sets. Pearson correlation between mean initial curvature and sulcus frequency was used to quantify their relationship.

### Statistical analysis

Each contour point was assigned to one of three curvature classes according to its arc segment in the initial trikyklon: high-curvature (small-radius arcs), medium-curvature, or low-curvature (large-radius arc). The mean probability of developing a sulcus was computed for each class and each simulation, yielding three distributions of 100 values. Differences between distributions were characterised using estimation statistics (`dabest` package; 5000 bootstrap samples, bias-corrected and accelerated 95% confidence intervals on unpaired mean differences).

## Acknowledgements

Funded by project DMOBE (ANR-21-CE45-0016), and the European Union’s Horizon 2020 research and innovation programme under the Marie Skłodowska-Curie grant agreement No101033485 (KH Individual Fellowship). For the purpose of open access, the author has applied a CC-BY public copyright licence to any Author Manuscript version arising from this submission. This work is supported by the ERC grant (UNFOLD, ERC-2023-SyG n°101118729). Funded by the European Union. Views and opinions expressed are however those of the author(s) only and do not necessarily reflect those of the European Union or the European Research Council Executive Agency. Neither the European Union nor the granting authority can be held responsible for them.

## References

Akula, S. K., Exposito-Alonso, D., & Walsh, C. A. (2023). Shaping the brain: The emergence of cortical structure and folding. Developmental Cell, 58(24), 2836–2849. 10.1016/j.devcel.2023.11.004

Alexander-Bloch, A. F., Shou, H., Liu, S., Satterthwaite, T. D., Glahn, D. C., Shinohara, R. T., Vandekar, S. N., & Raznahan, A. (2018). On testing for spatial correspondence between maps of human brain structure and function. NeuroImage, 178, 540–551. 10.1016/j.neuroimage.2018.05.070

Barnette, A. R., Neil, J. J., Kroenke, C. D., Griffith, J. L., Epstein, A. A., Bayly, P. V., Knutsen, A. K., & Inder, T. E. (2009). Characterization of Brain Development in the Ferret via MRI. Pediatric Research, 66(1), 80–84. 10.1203/pdr.0b013e3181a291d9

Bayly, P. V., Okamoto, R. J., Xu, G., Shi, Y., & Taber, L. A. (2013). A cortical folding model incorporating stress-dependent growth explains gyral wavelengths and stress patterns in the developing brain. Physical Biology, 10(1), 016005. 10.1088/1478-3975/10/1/016005

Budday, S., Steinmann, P., Goriely, A., & Kuhl, E. (2015). Size and curvature regulate pattern selection in the mammalian brain. Extreme Mechanics Letters, 4, 193–198. 10.1016/j.eml.2015.07.004

Caronia-Brown, G., Yoshida, M., Gulden, F., Assimacopoulos, S., & Grove, E. A. (2014). The cortical hem regulates the size and patterning of neocortex. Development, 141(14), 2855–2865. 10.1242/dev.106914

Chou, S.-J., Babot, Z., Leingärtner, A., Studer, M., Nakagawa, Y., & O’Leary, D. D. M. (2013). Geniculocortical Input Drives Genetic Distinctions Between Primary and Higher-Order Visual Areas. Science, 340(6137), 1239–1242. 10.1126/science.1232806

de Juan Romero, C., Bruder, C., Tomasello, U., Sanz-Anquela, J. M., & Borrell, V. (2015). Discrete domains of gene expression in germinal layers distinguish the development of gyrencephaly. The EMBO Journal, 34(14), 1859–1874. 10.15252/embj.201591176

Del-Valle-Anton, L., & Borrell, V. (2022). Folding brains: from development to disease modeling. Physiological Reviews, 102(2), 511–550. 10.1152/physrev.00016.2021

Foubet, O., Trejo, M., & Toro, R. (2019). Mechanical morphogenesis and the development of neocortical organisation. Cortex, 118, 315–326. 10.1016/j.cortex.2018.03.005

Fukuchi-Shimogori, T., & Grove, E. A. (2001). Neocortex Patterning by the Secreted Signaling Molecule FGF8. Science, 294(5544), 1071–1074. 10.1126/science.1064252

Garcia, K. E., Basinski, C., & Kroenke, C. D. (2025). Quantifying the timing of gyral and sulcal formation relative to growth in the ferret cerebral cortex. Developmental Neuroscience, 1–24. 10.1159/000544824

Garcia, K. E., Wang, X., & Kroenke, C. D. (2021). A model of tension-induced fiber growth predicts white matter organization during brain folding. Nature Communications, 12(1). 10.1038/s41467-021-26971-9

Garcia, K. E., Wang, X., Santiago, S. E., Bakshi, S., Barnes, A. P., & Kroenke, C. D. (2024). Longitudinal MRI of the developing ferret brain reveals regional variations in timing and rate of growth. Cerebral Cortex, 34(4). 10.1093/cercor/bhae172

Gilardi, C., & Kalebic, N. (2021). The Ferret as a Model System for Neocortex Development and Evolution. Frontiers in Cell and Developmental Biology, 9. 10.3389/fcell.2021.661759

Goriely, A. (2017). The Mathematics and Mechanics of Biological Growth. In Interdisciplinary Applied Mathematics. Springer New York. 10.1007/978-0-387-87710-5

Greiner, A., Kaessmair, S., & Budday, S. (2021). Physical aspects of cortical folding. Soft Matter, 17(5), 1210–1222. 10.1039/d0sm02209h

Heisenberg, C.-P., & Bellaïche, Y. (2013). Forces in Tissue Morphogenesis and Patterning. Cell, 153(5), 948–962. 10.1016/j.cell.2013.05.008

Heuer, K., & Toro, R. (2019). Role of mechanical morphogenesis in the development and evolution of the neocortex. Physics of Life Reviews, 31, 233–239. 10.1016/j.plrev.2019.01.012

Heuer, K., Gulban, O. F., Bazin, P.-L., Osoianu, A., Valabregue, R., Santin, M., Herbin, M., & Toro, R. (2019). Evolution of neocortical folding: A phylogenetic comparative analysis of MRI from 34 primate species. Cortex, 118, 275–291. 10.1016/j.cortex.2019.04.011

Heuer, K., Traut, N., de Sousa, A. A., Valk, S. L., Clavel, J., & Toro, R. (2023). Diversity and evolution of cerebellar folding in mammals. eLife, 12. 10.7554/elife.85907

Heuer, K., Traut, N., Aristide, L., Alavi, S. F., Herbin, M., Mars, R. B., Mylapalli, R., Najafipashaki, S., Sakai, T., Santin, M. D., Borrell, V., & Toro, R. (2025). Principles of neocortical organisation and behaviour in primates. openRxiv. 10.1101/2025.07.17.665410

Jablonka, E., & Lamb, M. (2020). Inheritance Systems and the Extended Synthesis. Cambridge University Press. 10.1017/9781108685412

Jia, F., Pearce, S. P., & Goriely, A. (2018). Curvature delays growth-induced wrinkling. Physical Review E, 98(3). 10.1103/physreve.98.033003

Krubitzer, L. (2007). The Magnificent Compromise: Cortical Field Evolution in Mammals. Neuron, 56(2), 201–208. 10.1016/j.neuron.2007.10.002

Lefèvre, J., & Mangin, J.-F. (2010). A Reaction-Diffusion Model of Human Brain Development. PLoS Computational Biology, 6(4), e1000749. 10.1371/journal.pcbi.1000749

Mallamaci, A., Muzio, L., Chan, C.-H., Parnavelas, J., & Boncinelli, E. (2000). Area identity shifts in the early cerebral cortex of Emx2−/− mutant mice. Nature Neuroscience, 3(7), 679–686. 10.1038/76630

Markello, R. D., Hansen, J. Y., Liu, Z.-Q., Bazinet, V., Shafiei, G., Suárez, L. E., Blostein, N., Seidlitz, J., Baillet, S., Satterthwaite, T. D., Chakravarty, M. M., Raznahan, A., & Misic, B. (2022). neuromaps: structural and functional interpretation of brain maps. Nature Methods, 19(11), 1472–1479. 10.1038/s41592-022-01625-w

Moffat, A., & Schuurmans, C. (2023). The Control of Cortical Folding: Multiple Mechanisms, Multiple Models. The Neuroscientist, 30(6), 704–722. 10.1177/10738584231190839

Namba, T., & Huttner, W. B. (2024). What Makes Us Human: Insights from the Evolution and Development of the Human Neocortex. Annual Review of Cell and Developmental Biology, 40(1), 427–452. 10.1146/annurev-cellbio-112122-032521

Nelson, C. M., Jean, R. P., Tan, J. L., Liu, W. F., Sniadecki, N. J., Spector, A. A., & Chen, C. S. (2005). Emergent patterns of growth controlled by multicellular form and mechanics. Proceedings of the National Academy of Sciences, 102(33), 11594–11599. 10.1073/pnas.0502575102

O’Leary, D. D. M. (n.d.). Do Cortical Areas Emerge from a Protocortex? In Brain Development and Cognition (pp. 217–230). Blackwell Publishers Ltd. 10.1002/9780470753507.ch12

O’Leary, D. D. M., & Nakagawa, Y. (2002). Patterning centers, regulatory genes and extrinsic mechanisms controlling arealization of the neocortex. Current Opinion in Neurobiology, 12(1), 14–25. 10.1016/s0959-4388(02)00285-4

Pang, J. C., Aquino, K. M., Oldehinkel, M., Robinson, P. A., Fulcher, B. D., Breakspear, M., & Fornito, A. (2023). Geometric constraints on human brain function. Nature, 618(7965), 566–574. 10.1038/s41586-023-06098-1

Pillai, E. K., & Franze, K. (2024). Mechanics in the nervous system: From development to disease. Neuron, 112(3), 342–361. 10.1016/j.neuron.2023.10.005

Rakic, P. (1988). Specification of Cerebral Cortical Areas. Science, 241(4862), 170–176. 10.1126/science.3291116

Ronan, L., & Fletcher, P. C. (2014). From genes to folds: a review of cortical gyrification theory. Brain Structure and Function, 220(5), 2475–2483. 10.1007/s00429-014-0961-z

Sawada, K., & Watanabe, M. (2012). Development of cerebral sulci and gyri in ferrets (Mustela putorius). Congenital Anomalies, 52(3), 168–175. 10.1111/j.1741-4520.2012.00372.x

Schwartz, E., Nenning, K.-H., Heuer, K., Jeffery, N., Bertrand, O. C., Toro, R., Kasprian, G., Prayer, D., & Langs, G. (2023). Evolution of cortical geometry and its link to function, behaviour and ecology. Nature Communications, 14(1). 10.1038/s41467-023-37574-x

Sharpe, J. (2019). Wolpert’s French Flag: what’s the problem? Development, 146(24). 10.1242/dev.185967

Shinmyo, Y., Hamabe-Horiike, T., Saito, K., & Kawasaki, H. (2022). Investigation of the Mechanisms Underlying the Development and Evolution of the Cerebral Cortex Using Gyrencephalic Ferrets. Frontiers in Cell and Developmental Biology, 10. 10.3389/fcell.2022.847159

Singh, A., Del-Valle-Anton, L., de Juan Romero, C., Zhang, Z., Ortuño, E. F., Mahesh, A., Espinós, A., Soler, R., Cárdenas, A., Fernández, V., Lusby, R., Tiwari, V. K., & Borrell, V. (2024). Gene regulatory landscape of cerebral cortex folding. Science Advances, 10(23). 10.1126/sciadv.adn1640

Subramanian, L., Remedios, R., Shetty, A., & Tole, S. (2009). Signals from the edges: The cortical hem and antihem in telencephalic development. Seminars in Cell & Developmental Biology, 20(6), 712–718. 10.1016/j.semcdb.2009.04.001

Tallinen, T., Chung, J. Y., Biggins, J. S., & Mahadevan, L. (2014). Gyrification from constrained cortical expansion. Proceedings of the National Academy of Sciences, 111(35), 12667–12672. 10.1073/pnas.1406015111

Tallinen, T., Chung, J. Y., Rousseau, F., Girard, N., Lefèvre, J., & Mahadevan, L. (2016). On the growth and form of cortical convolutions. Nature Physics, 12(6), 588–593. 10.1038/nphys3632

Tkačik, G., & Gregor, T. (2021). The many bits of positional information. Development, 148(2). 10.1242/dev.176065

Todd, P. H. (1982). A geometric model for the cortical folding pattern of simple folded brains. Journal of Theoretical Biology, 97(3), 529–538. 10.1016/0022-5193(82)90380-0

Toro, R. (2012). On the Possible Shapes of the Brain. Evolutionary Biology, 39(4), 600–612. 10.1007/s11692-012-9201-8

Toro, R., & Burnod, Y. (2005). A Morphogenetic Model for the Development of Cortical Convolutions. Cerebral Cortex, 15(12), 1900–1913. 10.1093/cercor/bhi068

Toro, R., & Heuer, K. (2025). Evolution of cortical folding. In Evolution of Nervous Systems. Elsevier. 10.1016/b978-0-443-27380-3.00039-7

Turing, A. M. (1952). The chemical basis of morphogenesis. Philosophical Transactions of the Royal Society of London. Series B, Biological Sciences, 237(641), 37–72. 10.1098/rstb.1952.0012

Tzika, A. C., Ullate-Agote, A., Zakany, S., Kummrow, M., & Milinkovitch, M. C. (2023). Somitic positional information guides self-organized patterning of snake scales. Science Advances, 9(24). 10.1126/sciadv.adf8834

Van Essen, D. C. (1997). A tension-based theory of morphogenesis and compact wiring in the central nervous system. Nature, 385(6614), 313–318. 10.1038/385313a0

Van Essen, D. C. (2020). A 2020 view of tension-based cortical morphogenesis. Proceedings of the National Academy of Sciences, 117(52), 32868–32879. 10.1073/pnas.2016830117

Vue, T. Y., Lee, M., Tan, Y. E., Werkhoven, Z., Wang, L., & Nakagawa, Y. (2013). Thalamic Control of Neocortical Area Formation in Mice. The Journal of Neuroscience, 33(19), 8442–8453. 10.1523/jneurosci.5786-12.2013

Welker, W. (1990). Why Does Cerebral Cortex Fissure and Fold? In Cerebral Cortex (pp. 3–136). Springer US. 10.1007/978-1-4615-3824-0_1

Wolpert, L. (1969). Positional information and the spatial pattern of cellular differentiation. Journal of Theoretical Biology, 25(1), 1–47. 10.1016/s0022-5193(69)80016-0

Xu, G., Knutsen, A. K., Dikranian, K., Kroenke, C. D., Bayly, P. V., & Taber, L. A. (2010). Axons Pull on the Brain, But Tension Does Not Drive Cortical Folding. Journal of Biomechanical Engineering, 132(7). 10.1115/1.4001683

Yoshino, M., Saito, K., Kawasaki, K., Horiike, T., Shinmyo, Y., & Kawasaki, H. (2020). The origin and development of subcortical U-fibers in gyrencephalic ferrets. Molecular Brain, 13(1). 10.1186/s13041-020-00575-8

